# p38 MAPK Signaling Enhances Reovirus Replication by Facilitating Efficient Virus Entry, Virus Capsid Uncoating and Post-Uncoating Steps

**DOI:** 10.1101/2022.04.20.488989

**Authors:** Adil Mohamed, Maya Shmulevitz

**Affiliations:** Department of Medical Microbiology and Immunology, Li Ka Shing Institute of Virology, University of Alberta, Edmonton, Alberta, Canada

## Abstract

Mammalian orthoreovirus serotype 3 Dearing an orphan human virus currently being pursued as an oncolytic virus in multiple phase I/II clinical trials. Previous clinical trials have emphasized the importance of patient pre-screening for prognostic markers to improve therapeutic success. However, currently, only generic cancer markers such as EGFR, Hras, Kras, Nras, Braf and/or p53 are utilized and have exhibited limited benefit in predicting therapeutic efficacy. Utilization of more specific markers that influence specific steps during reovirus replication could prove beneficial as prognostic markers. This study delineated the role of p38 MAPK signaling during reovirus infection and illustrated a connection between specific p38 MAPK isoforms and reovirus infection. Using a panel of specific p38 MAPK inhibitors and an inactive inhibitor analogue, we demonstrated that p38 MAPK signaling is essential for establishment of reovirus infection by enhancing reovirus endocytosis, facilitating efficient reovirus uncoating at the endo-lysosomal stage, and augmenting post uncoating replication steps. Using a broad panel of human breast cancer cell lines, we observed susceptibility of reovirus infection corresponded with virus binding and uncoating efficiency, which was strongly correlated with status of the p38β isoform. Together, our study proposes p38β as a potential prognostic marker for early stages of reovirus infection that are crucial to establishment of successful reovirus infection.

## INTRODUCTION

Mammalian orthoreovirus (reovirus) is a double-stranded RNA virus that is ubiquitous in nature and is non-pathogenic in humans. Reovirus type-3 Dearing PL strain ((T3D^PL^) (1)) is amongst the most clinically trialed oncolytic viruses and is currently being pursued as an oncolytic virotherapy in phase I-III clinical trials for breast cancer, gastro-intestinal cancer, and multiple myeloma (under clinical name Pelareorep) (2). In a randomized phase II trial of Pelareorep in metastatic breast cancer, median overall survival was extended from 10.4 months to 17.4, and despite no increase in response rate (P = 0.87) or progression-free survival (P = 0.87), received special protocol assessment agreement from FDA for a phase 3 clinical trial in metastatic breast cancer (3). Conversely in head and neck cancers, a phase III trial found reovirus therapy to provide no clinical benefit over standard of care (4). Importantly however, analysis of the patient cohorts suggested that inclusion of tumor prognostic markers such as EGFR, Hras, Kras, Nras, Braf and/or p53, signalling pathways known to enhance reovirus replication, slightly improved overall survival statistics, suggesting the immediate need for identification of additional prognostic markers (5–10). The current prognostic markers are upstream signaling modules that drive hundreds of downstream signaling cascades. For reovirus to initiate an infection, several steps have to be successfully performed; the virus must bind to the host cell, induce endocytosis and traffic to the lysosome, uncoat efficiently, initiate viral RNA transcription and establish viral factories. Current efforts are driven towards rational selection of prognostic markers that are important in each of these replication steps.

Upon virus exposure, the host cell responds by activating a myriad of signaling pathways in an attempt to inhibit establishment of the initial infection and/or restrict progeny virus spread to neighboring cells. Mitogen activated protein kinase (MAPK) stress signaling is activated upon exposure of the cell to viruses, growth factors and cytokines; and governs key cellular processes such as cell growth, apoptosis, differentiation, and inflammatory responses, which are important for successful virus establishment and spread. MAPK signaling is dictated by activation of three kinase families: Extracellular signal-regulated kinases (ERKs), c-Jun N-terminal kinases (JNKs) and p38 MAPKs (11, 12). Previous studies found that inhibitors of p38 MAPK and ERK signalling suppressed reovirus infection in cancer cells suggesting these pathways promote reovirus oncolysis (13). Mechanistically, ERK is essential for downregulating cellular interferon-mediated antiviral responses that support reovirus spread among cancer cell (10); but the molecular basis for reovirus-promoting effects of p38 MAPK signalling remain unknown.

In the context of cancer, p38 MAPK signaling is proposed to function as a dual regulator. In healthy tissues, p38 MAPK signaling maintains normal processes such as cell cycle, differentiation, and apoptosis. Contrarily, in cancer, p38 MAPK signaling functions to drive tumor progression by inducing processes such as migration, angiogenesis, and inflammation (11, 12). Replication of many viruses such as hepatitis B virus (HBV), hepatitis C virus (HCV), influenza virus and avian reovirus are dependent on p38 MAPK signaling (14–17). However, the precise mechanisms of action are distinct amongst the viruses. For example, p38 MAPK directly interacts with HCV, restricting virus assembly and replication, whereas p38 MAPK signaling expedites avian reovirus entry and endocytosis (14, 17).

In this study we sought to establish the role of p38 MAPK during reovirus infection, identify the potential mechanisms of action and determine the potential of utilizing p38 MAPK as a prognostic marker for reovirus oncolytic virotherapy. Using specific inhibitors and inactive analogs of p38 MAPK, we demonstrate that establishment of reovirus infection is dependent on p38 MAPK signaling. Specifically, reovirus endocytosis, trafficking to lysosomal compartments and post-uncoating steps are mediated by p38 MAPK signaling. Using a panel of breast cancer cell lines, we provide evidence that the p38β isoform correlates with the rate of reovirus uncoating.

## RESULTS

### Establishment of reovirus infection is dependent on p38 MAPK signaling

The p38 MAPK stress signaling pathway is induced during reovirus infection (10) and has been observed to play a beneficial role during initial reovirus infection (13) but the precise mechanisms of action are not well understood. Previous studies with reovirus and p38 MAPK signaling were performed in NIH/3T3 mouse fibroblast cells (13). Given that reovirus serotypes and laboratory strains induce different signaling cascades, we first sought to determine if our T3D laboratory strain (T3D^PL^) is capable of not only inducing activation of p38 MAPK, but also subsequent downstream target proteins in our model cell line, L929 mouse fibroblasts. The tumorigenic L929 cells are highly permissive to reovirus infection and commonly used to propagate and investigate reovirus mechanisms. Perturbations in p38 MAPK were performed using specific p38 MAPK inhibitors (SB202190, SB202580) or p38 MAPK activator (anisomycin). Many kinase-specific inhibitors are known have off-target interactions. To control for off-target effects of p38 MAPK inhibitors, we utilized SB202474, an analogue of SB202190 which shares similar chemical structure but is unable to interact with the p38 MAPK active site (18). Since the p38 MAPK signaling pathway is essential for cell survival, long term experiments using knockdown or knockout strategies were avoided.

Transformed cells can have dysfunctional signalling pathways due to acquired mutations, and therefore it was important to first determine if L929 cells maintain normal p38 MAPK signalling in absence of reovirus. To characterize the p38 MAPK signaling cascade, L929 cells were treated with anisomycin, a commonly used activator of p38 MAPK signalling, and Western blot analysis was performed for phospho-p38, its upstream kinase MKK3/6 and well-established downstream targets of p38; phospho-MSK1, phospho-MNK1, phospho-eEF2 and phospho-ATF2 (Figure 1A). As expected, anisomycin treatment increased P-MKK3/6, the upstream kinase of p38 MAPK and P-p38 (Figure 1B). Downstream of p38 MAPK, anisomycin triggered an increase in P-MSK1, a direct substrates of P-p38. The p38 signalling cascade seemed fully intact in L929 cells.

**FIG 1.**
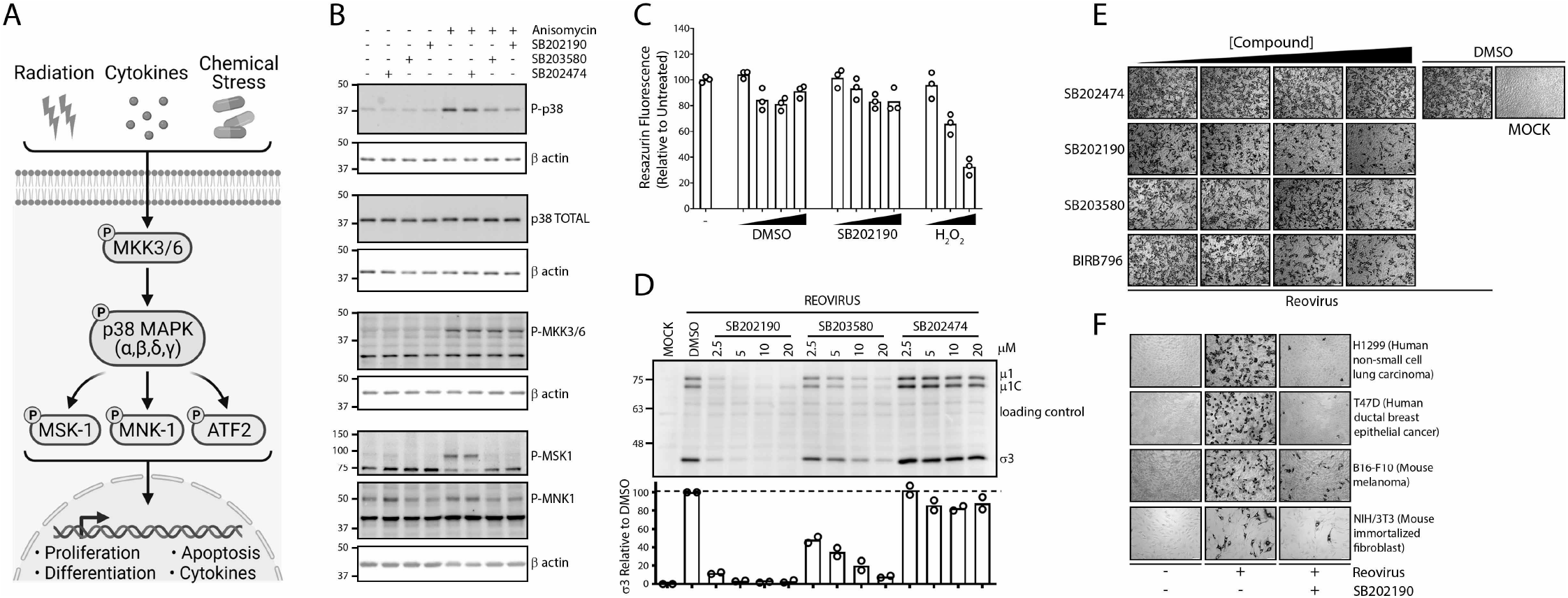
p38 MAPK signaling promotes establishment of reovirus infection. A) Summary of p38 MAPK activation and downstream signaling. B) L929 cells were treated with 10μM p38 MAPK inhibitors (SB2020190, SB203580) or inactive analog (SB202474) for 1hr followed by treatment with 500nM p38 MAPK activator (anisomycin) for 30min. Prototypic proteins along the p38 MAPK signaling pathway (A) were assessed using Western blot analysis. C) L929 cells were treated with various dilutions of SB202190 (20μM, 10μM, 5μM), respective DMSO for 12hrs, followed by a media change and additional incubation for 12hrs. Hydrogen peroxide (H_2_O_2_, 5mM, 1mM, 0.2mM) was incubated for 24hrs. Cell viability was assessed using resazurin fluorescence. (n=3) D and E) L929 cells were treated with indicated compound dilutions for 1hr at 37C. Cells were pre-chilled at 4C for 1hr, followed by reovirus addition and incubation at 4C for 1hr. Following extensive washing to remove unbound virus, media was replenished with indicated compound dilutions and incubated at 37C for 12hrs. D) Cell lysates were assessed for reovirus proteins using Western blot analysis. A background non-specific band was used as a loading control. E) Cells were fixed, and virus infected cells stained using reovirus-specific antibodies and BCIP/NBT substrate. F) Similar to E), except a panel of cell lines was used and SB2020190 was used at 10μM.

To establish the impact of p38 inhibitors, anisomycin-treated cells were subsequently also treated with p38 MAPK inhibitors (SB202190, SB203580) or inactive analogue (SB202474). P-MKK3/6 increased to a similar extent in anisomycin and p38 MAPK inhibitor treated cells, suggesting that inhibition of p38 MAPK signaling was not due to disruption in the upstream kinase of p38 MAPK. Although p38 MAPK inhibitors SB202190 and SB203580 decreased levels of phospho-p38 following anisomycin treatment, they did not fully eliminate p38 phosphorylation; this is typical among many other studies (19–24) and caused by continued phosphorylation of p38 MAPK by upstream kinase MKK3/6 (19, 20). In other words, the inhibitors bind to the ATP-binding active site of p38 and thereby stop phosphorylation of downstream targets such as MSK and MNK, but do not stop phosphorylation of p38 by upstream MKKs. Importantly, samples treated with p38 MAPK inhibitors in the presence of anisomycin resulted in reduction of P-MNK1, P-MSK1, corroborating inhibition of p38 MAPK signaling. The inactive analogue (SB202474) treated samples did not disrupt p38 MAPK signaling targets, supporting its use as an off-target control (Figure 1B).

Pharmacological treatments can sometimes cause toxicity to cells that produce confounding effects, so we assessed whether p38 MAPK inhibitors exhibited any toxicity at compound concentrations and treatment durations used in subsequent assays. L929 cells were treated with various doses of p38 MAPK inhibitor (SB202190), DMSO or hydrogen peroxide as a positive control for 12 hours, followed by removal of the drug/DMSO and further incubation for an additional 12 hours to permit ample time for cell death, if any, to occur. Cell viability was assessed using a resazurin dye fluorescence, an indirect measurement of cellular metabolism. At all concentrations of SB202190, no significant (>20%) toxicity was observed (Figure 1C). Hydrogen peroxide treatment demonstrated a dose-dependent decrease in cell viability with increasing concentration, confirming functionality of the resazurin assay. At all tested concentrations (5-40 μM) of SB202190 and DMSO, minimal cellular toxicity was observed, suggesting the safe use of 10-20μM SB202190 in our L929 cell line model.

Having validated functional p38 MAPK signaling in L929 cells and the specificity and safety of p38 MAPK inhibitors, these inhibitors were applied to evaluate the role of p38 MAPK in the context of reovirus infection. Cells were pre-treated with various concentrations of p38 MAPK inhibitors or the inactive analogue for 1hr prior to addition of reovirus. Cell lysates were collected at 12hpi and assessed using Western blot analysis for reovirus protein expression. Analysis of reovirus structural proteins μ1C and σ3, showed that treatment with p38 MAPK inhibitors markedly reduced reovirus protein expression relative to untreated and inactive analogue treated samples (Figure 1D). SB202190 and SB203580 reduced viral protein (μ1C and σ3) in a dose dependent manner, with SB202190 being the more potent inhibitor. Treatment at respective doses of SB202474 had minimal reduction in viral protein (μ1C and σ3), further confirming inhibitor specificity. In summary, the results suggest that basal p38 MAPK signaling is beneficial during the initial 12 hours of the reovirus infection cycle.

The reduced levels of reovirus proteins following p38 MAPK inhibitor treatment could reflect two possibilities: i) the number of cells infected by reovirus was reduced or ii) similar numbers of cells were infected but on a per-infected cell basis there was less reovirus replication. To distinguish between these possibilities and evaluate the effect of the p38 MAPK inhibitors on the proportion of reovirus infected cells, L929 cells were exposed to reovirus for 1hr and supplemented with media containing p38 MAPK inhibitors at various doses for 12hrs. Samples were fixed and reovirus-infected cells were visualized using immunocytochemistry. Reovirus infection in the untreated sample was similar to DMSO treated samples, suggesting that DMSO did not affect reovirus infection (Figure 1E). Both p38 MAPK inhibitors SB202190 and SB203580 reduced reovirus infection in a concentration dependent manner, with SB202190 exhibiting most-potent effects. Samples treated with the inactive p38 MAPK inhibitor analogue (SB202474) did not reduce reovirus infection, suggesting that off-target effects of SB203580 and SB202190 are not likely the cause of reovirus inhibition. Furthermore, a clinically-relevant p38 MAPK inhibitor BIRB796 also inhibited the number of cells productively infected by reovirus. Fewer reovirus-positive cells following p38 inhibitor treatments was also confirmed by flow cytometric analysis (data not shown). Altogether there was a clear reduction in the number of cells that became productively infected by reovirus when p38 MAPK activities were inhibited.

Finally, to establish whether p38 MAPK signaling was an essential pathway for reovirus infection in general rather than an L929-cell type dependent effect, reovirus infectivity was evaluated following p38 MAPK inhibitor (SB202190) treatment in mouse B16-F10 melanoma cells, human H1299 non-small cell lung carcinoma cells, human T47D breast ductal carcinoma cells and mouse NIH/3T3 non-transformed fibroblast cells. In all four cell lines tested, p38 MAPK inhibition drastically reduced the number of cells that became infected by reovirus (Figure 1F). Therefore, the p38 MAPK signaling pathway is important for establishment of initial reovirus infection, irrespective of cell type and transformation status.

### p38 MAPK signaling does not alter reovirus binding but expedites efficient viral entry

p38 MAPK signaling is important in numerous cellular processes such as endocytosis and endosomal trafficking. We hypothesized that treatment with p38 MAPK inhibitors prior to reovirus infection could potentially affect receptor expression on the cell surface and thereby reduce reovirus-cell binding and subsequent infection. To evaluate if p38 MAPK signaling affects cell attachment of reovirus, L929 cells were pretreated for 1hr with p38 MAPK inhibitor (SB202190), activator (anisomycin), or left untreated, followed by reovirus adsorption at 4°C to permit attachment but not entry. Unbound reovirus particles were removed by repeated rinsing and cell lysates were subjected to Western blot analysis for reovirus outer capsid proteins μ1C and σ3 (reflects cell-bound virus) or β actin (loading control). In both virus doses used, cell bound virus levels were unchanged between untreated and SB202190 or anisomycin treated samples (Figure 2A). As an alternative approach, attachment of S^35^ radiolabelled reovirus to L929 cells pre-treated with SB202190 was measured by liquid scintillation counting of total cell lysates. At all three reovirus dilutions, SB202190 treatment did not affect reovirus attachment (Figure 2B). Therefore, reduction in reovirus infection following inhibition of p38 MAPK signaling likely does not occur due to diminished reovirus-cell binding.

**FIG 2.**
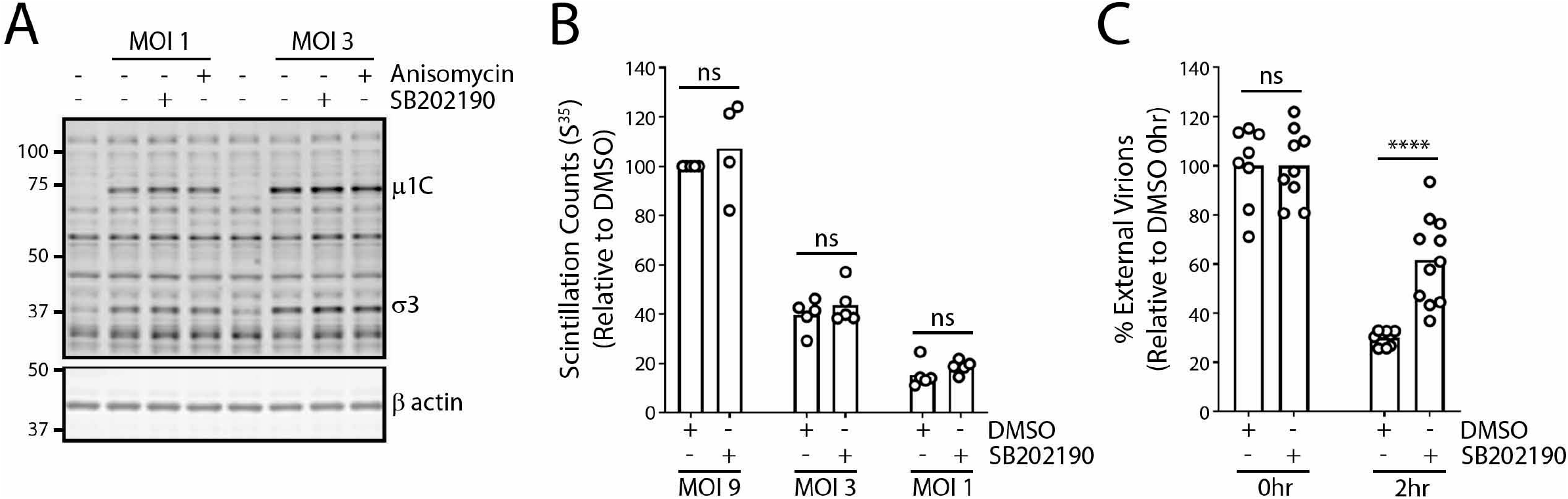
Reovirus entry but not binding is attenuated upon inhibition of p38 MAPK signaling. A) L929 cells were treated with indicated compounds (SB202190 10μM, anisomycin 500nM) for 1hr at 37C. Cells were pre-chilled at 4C for 1hr, followed by reovirus addition and incubation at 4C for 1hr. Following extensive washing to remove unbound virus cell lysates were collected and assessed for reovirus proteins using Western blot analysis. B) Cells were treated similar to A), except S^35^ radiolabelled reovirus was used, and cell lysates were collected and run on a scintillation counter. C) Cells were treated similar to A), except AlexaFluor 546 labeled reovirus was used. Cells were fixed at 0hr (immediately after binding) or after 2hr incubation at 37C. External virions were stained using reovirus specific antibodies without cell permeabilization. Cells were imaged using confocal microscopy and external (AlexaFluor 647) and internal (AlexaFluor 546) virions were quantified using the spot identification tool on Harmony High-Content Imaging and Analysis Software. Unpaired t-test * p < 0.05, ** p < 0.01, *** p < 0.001, **** p < 0.0001, ns p > 0.05.

Cellular entry of avian reovirus via dynamin and caveolin mediated endocytosis is p38 MAPK dependent (17). Since reovirus also shares these endocytosis pathways, we assessed whether reovirus entry also depends on p38 MAPK signaling. To quantitatively measure endocytosis, reovirus particles were covalently conjugated to Alexa Fluor™ 546 before addition to cells. Pre-conjugation of virus particles permitted quantification of both cell surface and internalized reovirus particles. To measure the percentage of virus particles that were not internalized after attachment to cells, extracellular virions were labelled on live cells immediately prior to fixation using anti-reovirus antibodies and respective secondary antibodies conjugated to Alexa Fluor™ 647. The percent of external virions (Alexa Fluor™ 647-positive) out of total (Alexa Fluor™ 546-positive) became a reflection of endocytosis. To measure the effects of p38 MAPK signalling on virus particle endocytosis, cells were pre-treated with p38 MAPK inhibitor (SB202190) for 1hr prior to exposure to Alexa Fluor™ 546 conjugated reovirus at 4°C for 1hr to permit virus binding. Samples were processed immediately after binding (0hpi) or following 2 hours of incubation at 37°C to permit endocytosis. Extracellular virions were then labelled using primary anti-reovirus and Alexa Fluor™ 647 secondary antibodies. Stained samples were fixed and imaged using confocal microscopy. At 0hpi, as expected, the average number of virus particles on the surface of cells was similar (Figure 2C). Conversely, at 2hpi, SB202190 treated samples had an average of 30% virion internalization, while untreated samples had an average of 62% virion internalization, suggesting that p38 MAPK signalling promotes efficient endocytosis of reovirus.

### Reovirus endosomal trafficking is facilitated by p38 MAPK signaling

Treatment with p38 MAPK inhibitor SB202190 caused 8 to 10-fold reduction in the number of reovirus-infected cells relative to untreated samples (Figure 1D), yet the inhibitor caused only a 2-fold reduction in virus internalization (Figure 2C); this suggested that a step downstream of endocytosis was also being inhibited during SB202190 treatment. While monitoring reovirus entry using Alexa Fluor™ 546 conjugated reovirus, we observed SB202190 treated samples had distinctly larger “spots” compared to untreated samples (Figure 3A), suggesting potential defects in endosomal trafficking.

**FIG 3.**
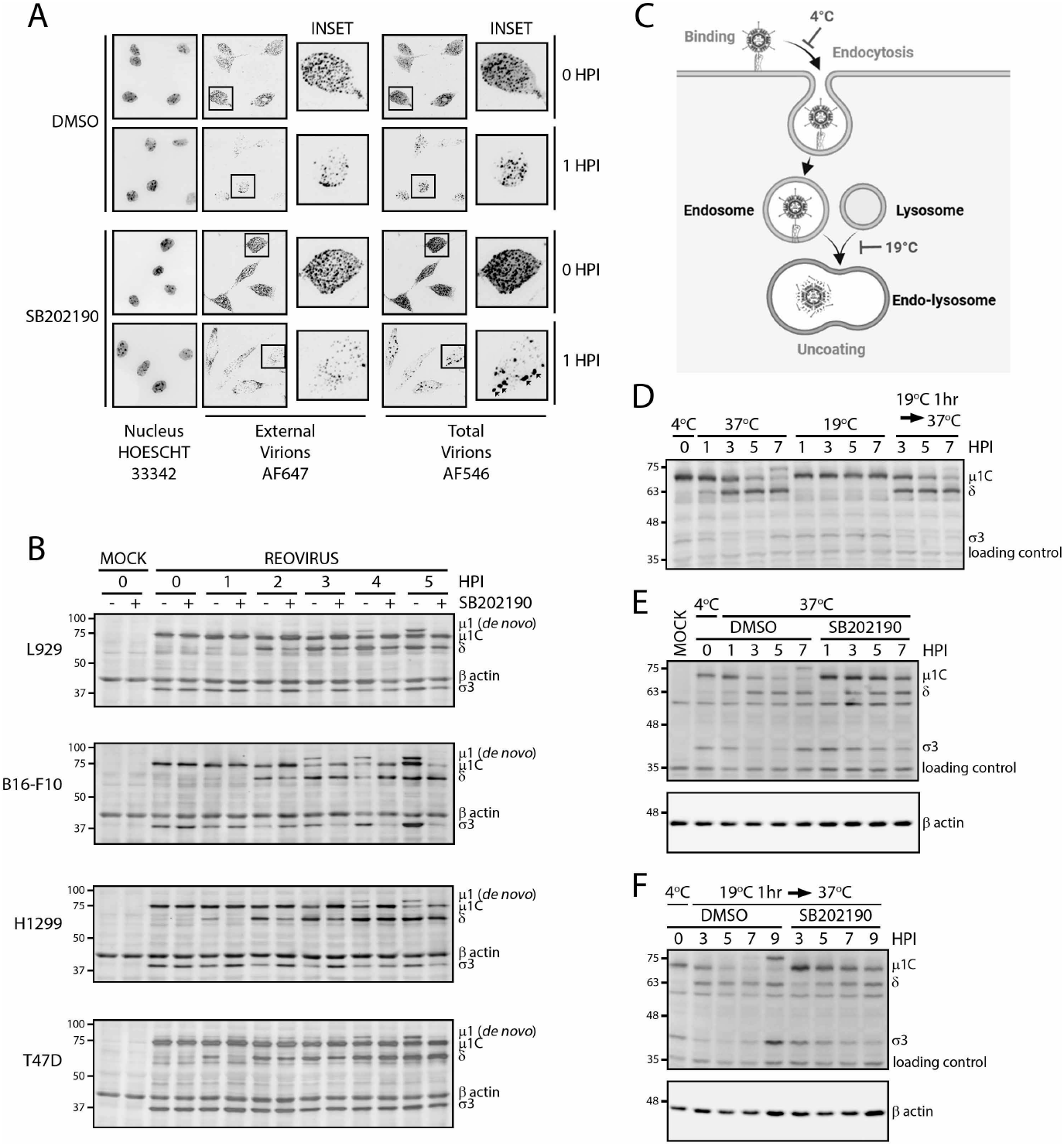
Reovirus uncoating is mediated by p38 MAPK signaling. A, B) Cells were treated with indicated compounds for 1hr at 37C. Cells were pre-chilled at 4C for 1hr, followed by reovirus addition and incubation at 4C for 1hr. Following extensive washing to remove unbound virus, media was replenished with indicated compounds and incubated at 37C. A) L929 cells were infected with AlexaFluor 546 labeled reovirus and cells were fixed at indicated timepoints followed by staining of external virions using reovirus specific antibodies and AlexaFluor 647. Nuclei were labelled using Hoescht 33342 cells were imaged using confocal microscopy. B, D-F) At indicated timepoints, cell lysates were collected and assessed for reovirus proteins using Western blot analysis. C) Summary of temperature effects on key steps of reovirus entry. D-F) Reovirus was bound to L929 cells, similar to A), and cells were incubated at indicated temperature in the presence of absence of SB202190. (SB202190 10μM).

Reovirus is composed of a core capsid that surrounds the dsRNA segmented genome, and is further encapsidated by an outercapsid composed of proteins σ1 (cell attachment), σ3 (outermost layer), and μ1 (intermediate protein between capsid and σ3). To initiate infection, the outercapsid protein σ3 must be proteolytically degraded and the underlying μ1 needs to be cleaved into a hydrophobic δ fragment to generate intermediate subviral particles (ISVPs) can penetrate cellular membranes and delivery of the core particle to the cytoplasm. During natural infection of cells in the gut, the outercapsid is degraded by digestive enzymes. But in cancer cells, for reovirus to establish an infection in the absence of digestive enzymes, reovirus-containing endocytic compartments must adjoin with lysosomes where proteolytic digestion of outer capsid proteins facilitates production of ISVPs that penetrate lysosome membranes to access the cytoplasm (25). We monitored reovirus capsid disassembly (uncoating) following SB202190 treatment in various cell lines (mouse L929 and B16-F10, human H1299 and T47D) using Western blot analysis for hallmarks of uncoating: degradation of σ3 and cleavage of μ1C to δ. At 0hpi, equivalent viral capsid proteins were observed between untreated and SB202190 treated samples, validating our previous observation that SB202190 does not alter virus-cell attachment (Figure 3B). In all cell lines tested, reovirus uncoating (μ1C to δ cleavage) was delayed in SB202190 treated samples, relative to untreated samples. Additionally, using *de-novo* μ1 as an indication of new viral protein synthesis, untreated samples had detectable *de-novo* μ1 as early as 3hpi, while no detectable *de-novo* μ1 was observed in SB202190 treated samples as late as 5hpi. Similar trends were observed in all cell lines tested. Therefore, p38 MAPK signaling corresponds with efficient reovirus uncoating, resulting in earlier initiation of viral protein synthesis.

To identify whether p38 MAPK signaling is important in endosome trafficking or lysosome fusion, we established a protocol to differentiate between these two possibilities. Early endosomes (pH 6.0-6.5) transition to late endosomes (pH 5.0-5.5) by means of a pH change mediated by the membrane traversing vacuolar ATPase (V-ATPase). Fusion of protease containing vesicles and additional activity of the V-ATPase facilitate the progression of the late endosome to an endo-lysosome (pH 4.5-5.0). In cells incubated at a low temperature (18-20°C), endocytosis and transition to late endosomes occur normally, but lysosome formation is inhibited, presumably due to vesicle fusion inhibition (26–29). Since lysosome formation is necessary for reovirus uncoating, we first determined if low temperature incubation (19°C) blocked reovirus uncoating (Figure 3C). L929 cells were bound by reovirus and reovirus uncoating was monitored at 0-7hpi using Western blot analysis for reovirus outer capsid proteins. At 37°C, reovirus uncoating proceeded with σ3 degradation and μ1C to δ cleavage. However, at 19°C σ3 degradation and μ1C cleavage was inhibited (Figure 3D). When the temperature was increased from 19°C to 37°C, reovirus progressed towards uncoating with the observation of σ3 degradation and μ1C to δ cleavage, suggesting a temperature-dependent reversible block in reovirus uncoating. Low (19°C) temperature incubation likely blocks reovirus uncoating by trapping virions in late endosomes incapable of accessing proteolytic lysosomal enzymes.

The ability to trap virions in uncoating-incapable late endosomes enabled us to assess if p38 MAPK signaling facilitates reovirus uncoating by promoting either late endosome-to-endo-lysosome progression or preceding endosome trafficking steps. Reovirus was bound to L929 cells and incubated at 37°C and/or 19°C with or without SB202190 in various combinations and processed using Western blot analysis to measure reovirus uncoating. Similar to our previous results, when maintained at 37°C, SB202190 treatment diminishes reovirus uncoating relative to untreated samples (Figure 3E). Conversely, samples were incubated at 19°C for 1hr to synchronously trap virus particles in endosomes prior to p38 MAPK inhibition. One hour was chosen because uncoating beings at 1hpi (Figure 3B). Cells were then treated with SB202190 for 30min prior to temperature shift to 37°C to initiate uncoating. Relative to DMSO treated samples, SB202190 treatment diminished reovirus uncoating even if added after the 19°C incubation (Figure 3F). In other words, even when endocytosis was enabled in absence of SB202190 treatment, the addition of SB202190 post-endocytosis still continued to prevent uncoating. Collectively, our results suggest that i) as expected, reovirus requires access to lysosomes for initiation and completion of outer capsid uncoating, and ii) p38 MAPK signaling facilitates efficient reovirus uncoating by modulating late endosome to lysosome transition.

### p38 MAPK signaling also independently enhances post-uncoating steps during reovirus replication

The p38 MAPK pathways affects many processes aside from vesicle transport, and it became important to establish if p38 MAPK effects on reovirus infection are solely a consequence of promoting virus entry or also other processes downstream of entry. For example, downstream substrates of p38 MAPK include factors involved in modulating protein translation (e.g MNK1/2 and MK2/3). Since reovirus utilizes the host translation machinery for viral protein synthesis, p38 MAPK signaling could impact the protein translation stage during reovirus infection. To evaluate the effects of p38 MAPK inhibition on post-entry steps, it became important to establish the timepoint at which virion uncoating was mostly completed; this would permit SB202190 treatment post-uncoating to evaluate effects on downstream steps of virus replication. Ammonium chloride (NH_4_Cl) is commonly used to neutralize lysosomal compartments, hence is capable of inhibiting reovirus uncoating (30). We added NH_4_Cl at various timepoints post infection and assessed reovirus infection at 12hpi using flow cytometry, reasoning that at the timepoint when the majority of uncoating was complete, NH_4_Cl would no long affect infection. Addition of NH_4_Cl at 0hpi had more than 90% inhibition on reovirus infection, which progressively increased with delayed addition of NH_4_Cl. Only a 2-fold reduction in reovirus infection was observed when NH_4_Cl was added at 2hpi. When NH_4_Cl was added at timepoints after 3hpi, minimal-to-no inhibition on reovirus infection was observed, suggesting that uncoating of virions essential for reovirus infection was accomplished between 3hpi and 4hpi (Figure 4A). Therefore, we could determine the effect of p38 MAPK signaling on post-reovirus uncoating steps by addition of SB202190 at timepoints after 3hpi.

**FIG 4.**
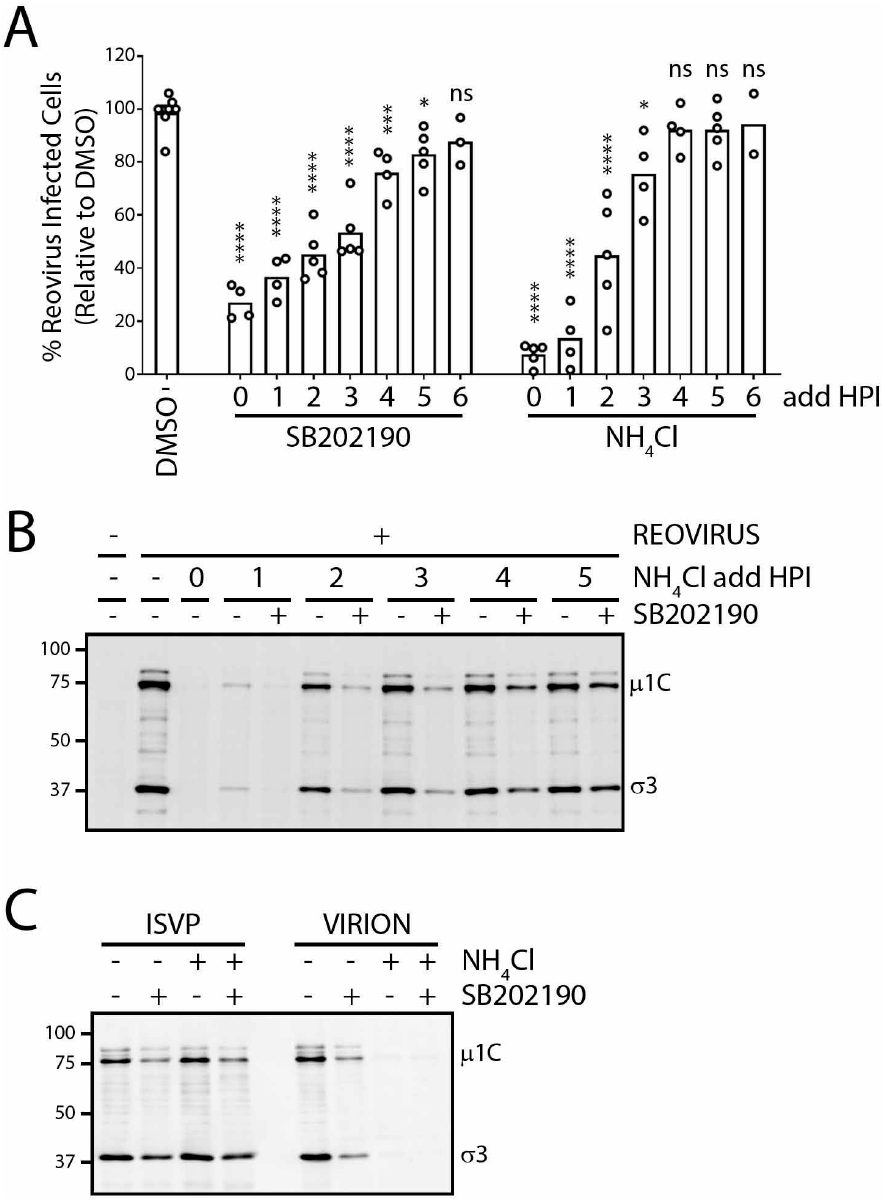
p38 MAPK signaling facilitates post-uncoating steps of reovirus replication. A) L929 cells were pre-chilled at 4C for 1hr, followed by reovirus addition and incubation at 4C for 1hr. Following extensive washing to remove unbound virus, media was replenished with and incubated at 37C. At indicated timepoints, compound was added, and cells were collected at 12HPI for flow cytometry to assess reovirus infected cells. B) Similar to A, except SB2020190 was added at 0hpi, whereas NH_4_Cl was added at indicated timepoints. Cell lysates were collected at 12hpi and assessed for reovirus proteins using Western blot analysis. C) Similar to A, except SB202190 and NH_4_Cl were added at 0hpi. Cell lysates were collected at 12hpi and assessed for reovirus proteins using Western blot analysis. (SB202190 20μM, NH_4_Cl 10mM). One-way analysis of variance with Dunnet’s multiple-comparison test to DMSO * p < 0.05, ** p < 0.01, *** p < 0.001, **** p < 0.0001, ns p > 0.05.

The levels of reovirus *de novo* protein expression at 12hpi was used to monitor the effects of p38 MAPK inhibition on post-entry steps. Specifically, L929 cells were infected with reovirus for different durations of time prior to addition of NH_4_Cl to stop further uncoating, plus/minus SB202190 at the same timepoint of NH_4_Cl addition to assess effects of p38 MAPK signalling on *de novo* virus protein expression at 12hpi by Western blot analysis. In agreement with the levels of reovirus infection observed by flow cytometry (Figure 4A), NH_4_Cl addition at ≥ 3hpi led to minimal effects on *de novo* virus protein expression since uncoating was mostly complete (Figure 4B). At every given timepoint, when NH_4_Cl and SB202190 were combined so that p38 MAKP inhibition only affects post-uncoating steps, there were still strong effects by SB202190 on *de novo* reovirus protein expression. As anticipated, addition of SB202190 at the onset on infection (1 and 2hpi) had more than 90% inhibition of viral proteins, suggesting that pre-reovirus uncoating steps are inhibited by SB202190. Importantly, when added at 3hpi or later, when reovirus uncoating was complete, reovirus protein synthesis was still inhibited by SB202190 by ≥50%. Accordingly, p38 MAPK signaling plays a role in post-reovirus uncoating steps that impact reovirus protein synthesis.

As an alternate strategy of circumventing the endocytic pathway during reovirus infection, we utilized reovirus ISVPs, that have been previously demonstrated to bypass the lysosomal pathway by either direct entry into the cytoplasm from the external cell membrane or cytoplasmic entry from the early endosomal vesicles (31). ISVPs consist of virions lacking outer capsid proteins σ3 and μ1 pre-cleaved to δ, and were generated by *in-vitro* treatment of reovirus particles to chymotrypsin. We reasoned that by infecting cells with reovirus ISVPs, the endocytosis and uncoating steps that are inhibited by SB202190 would be bypassed, and the effect of p38 MAPK signaling on post-uncoating steps of reovirus infection would be revealed. L929 cells were infected with reovirus ISVPs in the presence or absence of SB202190 treatment for 15hours, and reovirus proteins were monitored by Western blot analysis. NH_4_Cl treatment did not inhibit reovirus ISVP infection protein synthesis (Figure 4C), confirming that ISVP infection is lysosome-independent. In contrast, treatment with SB202190 inhibited *de novo* reovirus proteins synthesis even when cells were infected with ISVPs; this further suggested that post-uncoating reovirus replication steps are dependent on p38 MAPK signaling.

### Expression of p38β and p38δ MAPK isoforms correlate with reovirus uncoating in a breast cancer cell line panel

Given that p38 signalling affects multiple stages of reovirus replication, and p38 signalling status varies among cells, we wondered if there were relationships between the p38 signalling status of cells and the magnitude of reovirus infection. In reovirus clinical trials, patient responses are highly variable, and majority of patients fail to show clinical response relative to control test arm. However, with the inclusion of biomarkers such as EGFR, Hras, Kras, Nras, Braf and/or p53 mutations, patients expressing one or more of these biomarkers show an improved response to reovirus therapy compared to patents lacking any of the biomarkers (5–10). Given the importance of biomarkers in improving the prediction of patient response to reovirus therapy, we performed a screen to determine if p38 MAPK status correlates with reovirus infection in a panel of breast cancer cells (MDA-MB-468, MDA-MB-231, T-47D, Hs578T, BT-549, MCF7) from the NCI-60 cell line panel, characterizing reovirus infection versus various potential biomarkers including kinases of the p38 MAPK family.

To establish if p38 MAPK status relates with overall susceptibility of cancer cells to reovirus, and/or also to specific steps of reovirus infection affected by p38 MAPK signalling identified above, we first characterized the breast cancer cell panel with respect to overall susceptibility, entry, and post-entry steps of replication. To measure overall susceptibility of the breast cancer cell line to reovirus infection, reovirus infection was allowed to proceed for 12-15hrs and the frequency of productively-infected cells was assessed using immunocytochemistry. The dose of reovirus required to achieve approximately 50% infected cells was determined for each cell type. Note that multiplicity of infection (MOI) was calculated based on reovirus titers measured using highly susceptible L929 cells; by Poisson distribution, a cell line equally susceptible to reovirus as L929 cells should therefore show 63.2% of cells infected at an MOI of 1. Accordingly, MDA-MB-468 and T-47D cells were highly susceptible similar to L929 cells since multiplicities of 0.3-1 were sufficient to cause ≥50% infected cells (Figure 5A). MDA-MB231 and MCF7 cells were intermediate, requiring MOI 20-60 for ≥50% infected cells. Hs578T and BT-549 cells were highly resistant to initial reovirus infection, requiring ≥10 times higher MOIs than MDA-MB231 and MCF7 cells for ≥50%. Relative to the other breast cancer cell lines in our panel, BT-549 cells had a long doubling time (~60hrs) and was therefore excluded from subsequent analysis to eliminate the confounding effects of cell division to eliminate doubling time as a confounding variable.

**FIG 5.**
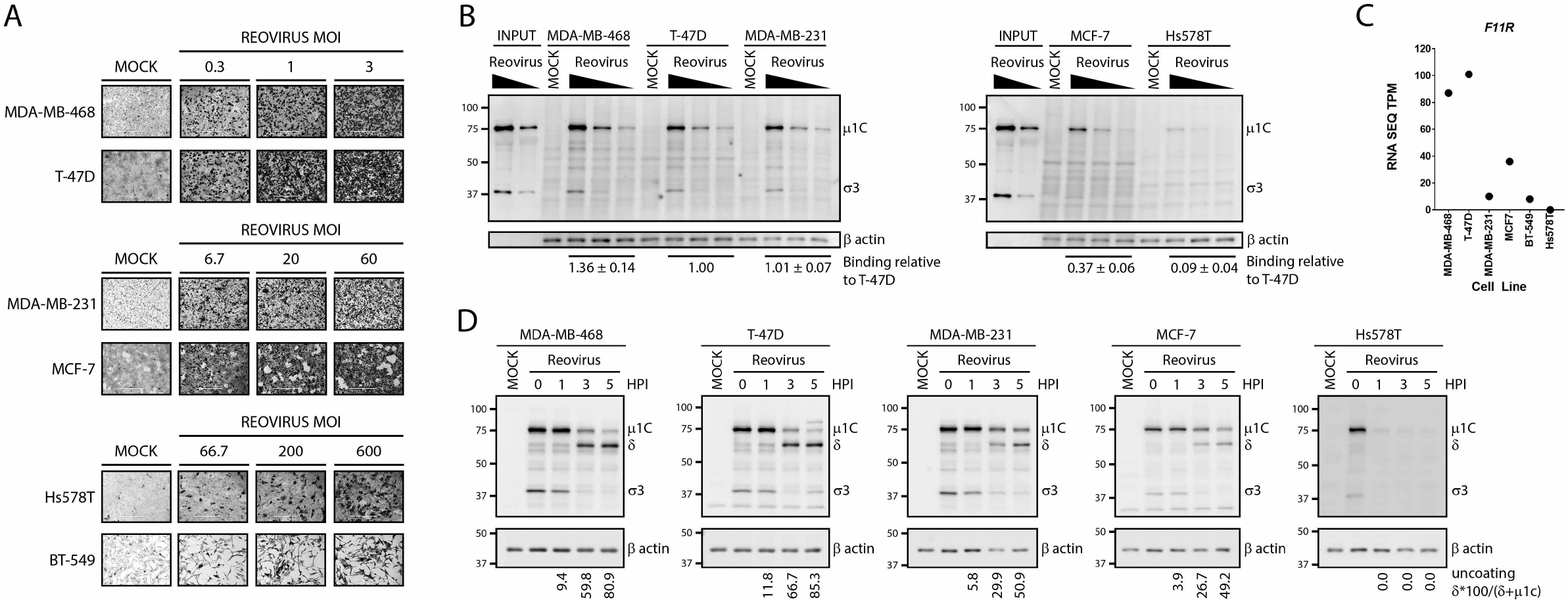
Differential susceptibility of breast cancer cell lines to reovirus infection. A) Breast cancer cell lines were infected with indicated doses of reovirus for 12hrs. Cells were fixed, and virus infected cells stained using reovirus-specific antibodies and BCIP/NBT substrate. MOI were calculated as per titres on L929 cells. B, D) Breast cancer cell lines were pre-chilled at 4C for 1hr, followed by reovirus addition and incubation at 4C for 1hr. Following extensive washing to remove unbound virus, cell lysates were collected at B) 0hpi, or D) indicated timepoints after 37C incubation and assessed for reovirus proteins using Western blot analysis. B) Input indicates virus inoculum. C) Basal transcript levels of F11R were obtained from the EMBL-EBI Expression Atlas database (https://www.ebi.ac.uk/gxa/home), and compared between cell lines.

Reovirus binds cells via cell surface junction adhesion molecule (JAM-A) and sialic acids (SA), and since cells can vary in their JAM and SA expression levels, reovirus binding was evaluated as a potential step for differential infection among breast cancer cells. Cells were exposed to reovirus at 4°C for 1hr, rinsed extensively, and subjected to Western blot analysis to measure levels of bound reovirus structural proteins. For each cell line, β-actin was normalized on a per cell basis and used as a correction factor to account for differences in cell size and hence plate seeding, between the cell lines. Reovirus exhibited strong attachment to the highly susceptible MDA-MB-468 and T47D cells and the moderately-susceptible MDA-MB-231 cells, but also ~40% attachment to the moderately susceptible MCF7 cell line (Figure 5B). There was very low attachment to Hs578T cells, suggesting that resistance of Hs578T to reovirus infection is likely due to inability of the virus to bind to these cells. Using the European Molecular Biology Laboratory (EMBL) cancer cell line Expression Atlas database, we extracted expression data for F11R (JAM-A), the primary reovirus cell binding receptor. While all the other cell lines had F11R expression varying from 10 to 101, Hs578T had zero expression (Figure 5C, F11R), suggesting that lack of JAM-A on Hs578T might contribute to poor reovirus binding and subsequent establishment of infection.

Since reovirus uncoating is a rate-limiting step during establishment of infection, we assessed whether reovirus uncoating levels correspond with reovirus susceptibility in our cell line panel. Reovirus uncoating was monitored by quantifying μ1C to δ cleavage at various timepoints post reovirus binding using Western blot analysis. For Hs578T cells, even though low levels of reovirus were detected immediately after binding (0hpi), no viral protein was detected at subsequent timepoints (1-5hpi), suggesting that the few bound virions had detached from the cell and were not internalized (Figure 5D). Reovirus uncoating in the highly susceptible MDA-MB-468 and T-47D cells occurred most-rapidly, reaching ~60% uncoating by 3hpi and >80% uncoating by 5hpi (Figure 5C). The moderately susceptible MDA-MB231 and MCF-7 cell lines supported intermediate reovirus uncoating rates, achieving ~30% uncoating by 3hpi and ~50% uncoating by 5hpi. The relative rates of uncoating correlated well with the infectivity assessed. The correlation between susceptibility and uncoating was then validated on several additional cell lines (Figure 6A, B). The H1299 human lung carcinoma cell line was highly susceptible (equivalent to L929, MDA-MB-468 and T-47D cells) and showed rapid uncoating (~60% uncoating by 3hpi). HCT116 human colorectal carcinoma cells were moderately susceptible (similar to MDA-MB-231 and MCF7) and exhibited ~50% uncoating by 3hpi (Figure 6B). The overall trend suggested that given the ability of reovirus to attached to breast cancer cells, the rate of reovirus uncoating correlates with overall reovirus infection.

**FIG 6.**
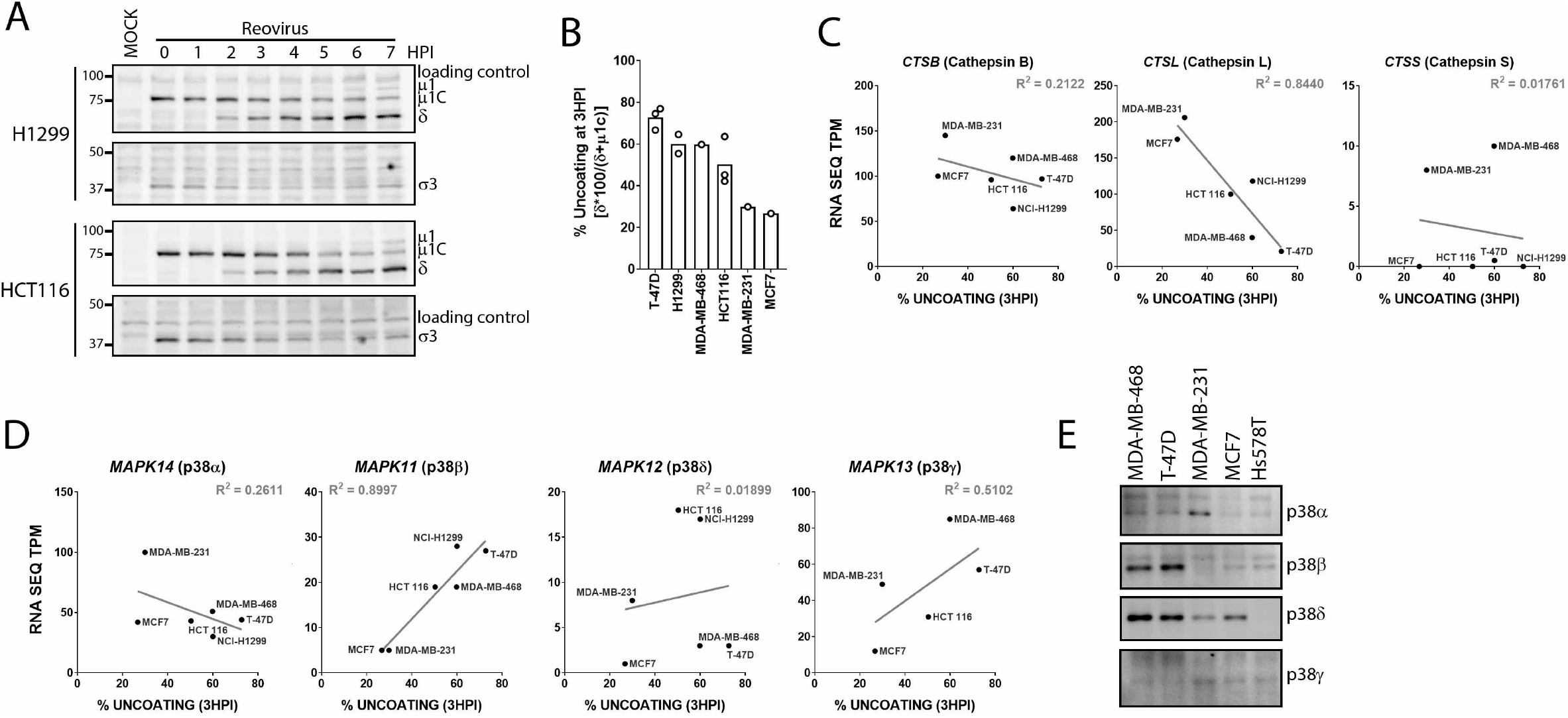
Reovirus uncoating correlates with expression of p38β isoform. A) H1299 and HCT116 cell lines were pre-chilled at 4C for 1hr, followed by reovirus addition and incubation at 4C for 1hr. Following extensive washing to remove unbound virus, cell lysates were collected at indicated timepoints after 37C incubation and assessed for reovirus proteins using Western blot analysis. B) Quantification of % capsid uncoating at 3HPI. C, D) Basal transcript levels of C) CTSB, CTSL, CTSL, D) MAPK14, MAPK11, MAPK12, MAPK13 were obtained from the EMBL-EBI Expression Atlas database (https://www.ebi.ac.uk/gxa/home), and compared between cell lines (A, B) or against % uncoating at 3HPI (C, D). Indicated cell lines were normalized to 1×106cells and equal volume of cell lysates were processed by Western blot analysis for p38 MAPK isoforms.

Intracellular uncoating of reovirus was previously shown to be mediated by lysosomal cathepsins B, L, and S (32). The obvious question was therefore whether differential rates of uncoating among cell lines was correlated to and levels of cathepsin. The NCI-60 cancer cell panel has been extensively characterized with respect to cellular mRNA expression levels and publicly available through the EMBL-EBI Expression Atlas database. When levels of each cathepsin transcript were plotted against percentage of reovirus uncoating at 3hpi for each cell line, there was small positive correlation with cathepsin B (R^2^=0.2122) and a strong negative correlation with cathepsin L (R^2^=0.8440), but poor correlation with cathepsin S (R^2^=0.01761) (Figure 6C). Levels of cathepsin L transcripts may therefore be a inverse prognostic marker of reovirus uncoating.

Having determined that p38 MAPK signalling was required for efficient steps leading to uncoating, we next assessed if uncoating rates also correlated with p38 MAPK isoform levels. Using the EMBL-EBI Expression Atlas database, the transcript levels of each p38 MAPK isoform were plotted against percentage of reovirus uncoating at 3hpi for each cell line. Hs578T and BT-549 were excluded from the analysis since reovirus uncoating was not determined due to low binding or slow cell growth/division, respectively. Correlation analysis by linear regression was then used to determine if p38 MAPK isoform expression correlated with reovirus uncoating efficiency. Of the four p38 MAPK isoforms, MAPK14 (encodes p38α) and MAPK12 (encodes p38δ) has poor correlation to reovirus uncoating (Figure 6D). The expression of MAPK13 (encodes p38γ) is tissue dependent with very low to no expression in lung tissue. Excluding H1299 lung cancer cell lines, MAPK13 (encodes p38γ) had moderate (R^2^ = 0.5102) correlation to reovirus uncoating. On the other hand, MAPK11 (encodes p38β) demonstrated a very strong (R^2^ = 0.8997) correlation to reovirus uncoating (Figure 6D). Finally, to further validate p38 MAPK isoform expression in the breast cancer cell lines, protein levels of p38 MAPK isoforms were determined by Western blot analysis using isoform-specific antibodies. All cell lines were standardized to equivalent number of cells per μl of lysate to account for differences in cell size. Similar to RNA levels, protein expression of p38β and p38δ were highly expressed in MDA-MB-468 and T-47D; the two cell lines with the fastest reovirus uncoating (Figure 6E). Together with the correlation analysis, the results suggests that the MAPK11 gene and its p38β protein isoform offer the strongest association with efficiency of reovirus uncoating and infection.

## DISCUSSION

Ongoing clinical trials with Pelareorep include pre-screening patients for activation and/or overexpression mutations in EGFR, Hras, Kras, Nras, Braf and/or p53 genes. However, the inclusion of these prognostic markers has shown very limited success, primarily due to these biomarkers being the most common occurring mutations in cancers. Additionally, since the EGFR and its RAS signaling cascade triggers a wide array of downstream cascades, an aberrant cancer inducing mutation in a downstream pathway will nullify the use of EGFR/RAS as prognostic markers. Therefore, there is a need to identify novel downstream prognostic markers associated with enhanced reovirus infection. In this study, we identify p38 MAPK signaling as an essential pathway for establishment of reovirus infection, specifically during reovirus endocytosis, lysosomal uncoating and post uncoating steps. Moreover, using a breast cancer cell line panel, we demonstrate that reovirus uncoating correlates with expression of p38β.

During reovirus infection, trafficking of the virion to the lysosome is a rate limiting step; virions in early endosomes are unable to initiate infection, while virions in lysosomes for an extended duration are inactivated (33). Our results suggest that p38 MAPK signaling mediates internalized reovirus vesicular trafficking to lysosomes; specifically finding that p38 inhibitor treatment restricts intracellular reovirus uncoating, internalized virions reside in larger vesicles following p38 inhibitor treatment, and impedance of late endosome and lysosome fusion restricts reovirus uncoating in p38 inhibitor treated cells. The p38 MAPK signaling pathway is involved in various aspects of endosome trafficking. Endosomes are classified in accordance with the presence of specific markers; Rab5 and early endosome antigen 1 (EEA1) for early endosomes, Rab7 and Rab9 for late endosome (34, 35). It was demonstrated that phosphorylation of guanine nucleotide dissociation inhibitor (GDI) by p38 MAPK results in formation of cytoplasmic P-GDI-Rab5 complexes which are required for Rab5 delivery, docking and loading onto early endosome membranes. Artificial activation of p38 MAPK using H_2_O_2_ or UV treatment increased internalization rates and early endosome formation, whereas inhibition of p38 MAPK using SB203580 had the opposite phenotype of decreased internalization rates and early endosome formation (36, 37). Therefore, our observation that p38 MAPK signaling modulates reovirus internalization could potentially be related to p38 MAPK-GDI-Rab5 mediated early endosome formation. Moreover, our result of internalized fluorescent virions forming larger vesicles in the presence of a p38 MAPK inhibitor, could also be an indication of defective early endosome formation.

Extrusion of Rab5 from early endosome membranes occurs concurrently with loading of Rab7 resulting in the transition to late endosomes. Rab7 containing late endosomes are the site of lysosome fusion and subsequent degradation of endosome contents (34, 38). In our temperature controlled endosomal trafficking experiment, we established that incubation at 19°C allows endocytosis, however virus uncoating fails to occur, indicating a block in access to lysosomal contents. When reovirus infected cells at 19°C were moved to 37°C, reovirus uncoating progressed, suggesting successful interaction of virions with lysosomal contents. Interestingly, in the presence of a p38 MAPK inhibitor, reovirus uncoating did not occur when moved from 19°C to 37°C, proposing a p38 MAPK regulated defect in either late endosome/lysosome fusion and/or lysosome biogenesis. Indeed, p38 MAPK was demonstrated to phosphorylate LAMP2A bound to lysosomes resulting in dysfunctional lysosomal biogenesis (39).

Establishment of a virus infection in cells relies on the success of multiple steps; binding, entry, viral genome replication, viral protein synthesis, virion assembly and virus release. Hence, successful identification of prognostic markers requires in-depth understanding of host signaling pathways important during each stage of the virus replication cycle. In a panel of breast cancer cell lines, we quantified reovirus binding and uncoating and attempted to correlate previously established factors important in these steps. Reovirus binding to cells occurs via cell surface JAM-A (F11R) and sialic acids, whereas endocytosis is triggered by interaction with β1 integrin (40–42). In our breast cancer cell line panel analysis, we observed that any expression of F11R albeit low or high, was sufficient for reovirus binding. However, lack of F11R expression abrogated reovirus binding, suggesting that F11R could be utilized as a potential tumor biomarker to assess for reovirus binding. In a study performed by Kelly, K.R *et al*. in a panel of advanced multiple myeloma cell lines, they also observed that lack of F11R (JAM-A) restricted reovirus infection, further supporting the use of F11R (JAM-A) as a prognostic marker for reovirus therapy (43). Reovirus uncoating is modulated by lysosomal cathepsins (32), however none of the cathepsins (CTSB, CTSS, CTSL) were positively correlated with the rate of reovirus uncoating, at least at the transcript level. Strikingly, we observed a very strong positive correlation between MAPK11 (p38β) and the rate of reovirus uncoating, upholding our evidence of the importance of p38 MAPK signaling in reovirus entry and uncoating.

In summary, our study elucidates the functional importance of p38 MAPK signaling during reovirus entry, uncoating and post-uncoating steps during infection and identifies the p38β isoform as a potential biomarker for reovirus uncoating and overall susceptibility.

## MATERIALS AND METHODS

### Cell lines

All cell lines were grown at 37°C at 5% CO2 and all media was supplemented with 1x antibiotic antimycotics (A5955, Millipore Sigma). Except for NIH/3T3 media that was supplemented with 10% NCS (N4637, Millipore Sigma), all other media was supplemented with 10% FBS (F1051, Millipore Sigma). L929, NIH/3T3, H1299, and B16-F10 cell lines (Dr. Patrick Lee, Dalhousie University) were generous gifts. BT-549, Hs578T, MCF7, T-47D, MDA-MB-231 and MDA-MB468 cell lines were purchased from ATCC as part of the NCI-60 cell line panel. L929 cell line was cultured in MEM (M4655, Millipore Sigma) supplemented with 1× non-essential amino acids (M7145, Millipore Sigma) and 1mM sodium pyruvate (S8636, Millipore Sigma). L929 cell line in suspension were cultured in Joklik’s modified MEM (pH 7.2) (M0518, Millipore Sigma) supplemented with 2g/L sodium bicarbonate (BP328, Fisher Scientific), 1.2g/L HEPES (BP310, Fisher Scientific), 1× non-essential amino acids (M7145, Millipore Sigma) and 1mM sodium pyruvate (S8636, Millipore Sigma). H1299, BT-549, Hs578T, MCF7, T-47D, MDA-MB-231 and MDA-MB468 cell lines were cultured in RPMI (R8758, Millipore Sigma). NIH/3T3 and B16-F10 cell lines were cultured in DMEM (D5796, Millipore Sigma) supplemented with 1mM sodium pyruvate (S8636, Millipore Sigma). All cell lines were routinely assessed for mycoplasma contamination using 0.5μg/ml Hoechst 33352 (H1399, ThermoFisher Scientific) staining or PCR (G238, ABM).

### Antibodies

Rabbit anti-reovirus pAb (1:10,000) from Dr. Patrick Lee (Dalhousie University), Rabbit anti P-p38 (1:1,000, CST Cat #9215S), Rabbit anti p38 (1:500, SCBT Cat#SC-535), Mouse anti-β actin (1:1000, SCBT Cat#47778), Rabbit anti P-ATF2 (1:1,000, CST Cat# 5112S), Rabbit anti P-MKK3/6 (1:1,000, CST Cat#9231S), Rabbit anti P-MSK1 (1:1,000, CST Cat# 9595P), Rabbit anti P-MNK1 (1:1,000, CST Cat# 2111S), Rabbit anti p38α (1:1,000, CST Cat# 9218), Rabbit anti p38β (1:1,000, CST Cat# 2339), Rabbit anti p38δ (1:1,000, CST Cat#2308), Rabbit anti p38γ (1:1,000, CST Cat#2307), Goat anti rabbit HRP (1:10,000, JIR Cat# 111-035-144), Goat anti mouse HRP (1:10,000, JIR Cat# 115-035-146, Goat anti-rabbit AP (1:10,000, JIR Cat# 111-055-144), Goat anti rabbit Alexa Fluor 647 (1:2,000, JIR Cat# 111-605-144), Goat anti-rabbit Alexa Fluor 488 (1:2,000, JIR Cat# 111-545-144), Goat anti-mouse Alexa Fluor 647 (1:2,000, JIR Cat# 115-605-146). CST = Cell Signaling Technology, SCBT = Santa Cruz Biotechnology, JIR = Jackson ImmunoReaseach.

### Compounds

SB202190 (100mM in DMSO, MilliporeSigma Cat# S7067), SB203580 (50mM in DMSO, MilliporeSigma Cat# S8307), SB202474 (50mM in DMSO, MilliporeSigma Cat# 559387), Anisomycin (18.85mM in DMSO, MilliporeSigma Cat# A9789), BIRB796 (125.6mM in DMSO, MilliporeSigma Cat# 506172), Ammonium chloride (NH_4_Cl) (4M in H2O, MilliporeSigma Cat# A9434), Hydrogen peroxide (H_2_O_2_) (100mM in H2O, FisherScientific Cat# H312-500).

### Reovirus stocks

Seed stock lysates were plaque purified and second or third passage L929 cell lysates were used as spinner culture inoculums. Cesium chloride purified preparations of reovirus were performed as previously described in detail (1).

### Resazurin Cell Viability Assay

Resazurin stock solution was prepared at 440μM in PBS and stored in aliquots at 4°C. Immediately prior to addition to wells, resazurin stock solution was dilution 1/10 in PBS and 10ul was added per 96 well. Samples were read on plate reader (FLUOstar Optima, BMG Lab Tech) every 30min for 2hrs, until signal saturation. Fluorescence was measured at excitation 520nm an emission 584nm.

### Radiolabeled (S35) reovirus virions

At 9hr post reovirus infection in L929 cells, cells were washed and media replaced with with methionine/cysteine free media supplemented with 35S-methionine at 100μCi/ml and dialyzed FBS at 10% final. At 20hpi, cells were washed with PBS and lysed in RIPA (50mM Tris pH 7.4, 150mM NaCl, 1% IGEPAL CA-630 (NP-40), 0.5% sodium deoxycholate) supplemented with protease inhibitor cocktail (11873580001, Roche). Lysate was layered on a 20% sucrose cushion and centrifuged at 100,000g for 90min at 4°C. Pellet containing radiolabeled reovirus was resuspended in PBS and stored at 4°C.

### Fluorescent (Alexa Fluor 546) labeled reovirus virions

Reovirus stock was diluted to 3×10^12^ particles in freshly made 0.05M sodium bicarbonate (pH 8.5) and Alexa Fluor 546 dye was added to 12μM final dye concentration. After incubation at 4°C for 90min, an overnight dialysis (MWCO 10-20KDa) in PBS was performed and virus was collected and stored at 4°C.

### Western blot analysis

Cells were washed with PBS and lysed in RIPA buffer (50mM Tris pH 7.4, 150mM NaCl, 1% IGEPAL CA-630 (NP-40), 0.5% sodium deoxycholate) supplemented with protease inhibitor cocktail (11873580001, Roche) and phosphatase inhibitors (1mM sodium orthovanadate, 10mM β-glycerophosphate, 50mM sodium fluoride). Following addition of 5x protein sample buffer (250mM Tris pH 6.8, 5% SDS, 45% glycerol, 9% β-mercaptoethanol, 0.01% bromophenol blue) for a final 1x protein sample buffer, samples were heated for 5min at 100°C and loaded onto SDS-acrylamide gels. After SDS-PAGE, separated proteins were transferred onto nitrocellulose membranes using the Trans-Blot^®^ Turbo™ Transfer System (Bio-Rad). Membranes were blocked with 3% BSA/TBS-T (blocking buffer) and incubated with primary and secondary antibodies, with 3x wash steps between antibody incubation steps. Membranes with HRP-conjugated antibodies were exposed to ECL Plus Western Blotting Substrate (32132, ThermoFisher Scientific). Membranes were visualized using ImageQuant LAS4010 imager (GE Healthcare Life Sciences) and densitometric analysis was performed by using ImageQuant TL software (GE Healthcare Life Sciences).

### Flow cytometry

Cells were detached with trypsin and washed with media containing FBS to quench trypsin. Cells were fixed with 4% paraformaldehyde 4°C for 30min, followed by permeabilization and blocking in 3% BSA/PBS/TritonX-100 at room temperature for 1hr. Samples were spiked with primary antibody (rabbit anti-reovirus pAb, 1:10,000) and incubated overnight at 4°C. Samples were washed twice and incubated with secondary antibody (goat anti-rabbit Alexa Fluor 647, 1:2,000 dilution) and incubated at room temperature for 1hr. Samples were washed twice and analyzed using FACSCanto (BD Biosciences). Flow cytometry data was analyzed using FSC Express 5 (De Novo Software). A minimum of 10,000 total cells were collected for each sample.

## ACKNOWLEDGMENTS

This publication is supported through project grants to MS from the Canadian Institutes of Health Research (CIHR), the Cancer Research Society (CRS), the Canadian Cancer Society Research Institute (CCSRI). A salary award to MS from the Canada Research Chairs (CRC) and infrastructure support to MS from the Canada Foundation for Innovation (CFI). A.M. received scholarships from an Alberta Cancer Foundation Graduate Studentship, a University of Alberta Faculty of Medicine and Dentistry/Alberta Health Services Graduate Recruitment Studentship, and a University of Alberta Doctoral Recruitment Award.

